# Nutrient-limited conditions reveal the activity of a minor groove binder, MGB-BP-3, against *Escherichia coli*

**DOI:** 10.64898/2025.12.22.695926

**Authors:** Kaveh Eskandari, Louise C. Young, Jessica M A Blair, Katarina Oravcova, Fraser J. Scott

## Abstract

**Background:** Antimicrobial resistance demands new antibiotics, but conventional screening in nutrient-rich media often fails to predict *in vivo* efficacy, especially against Gram-negative bacteria. Minor groove binders (MGB), such as MGB-BP-3, are a promising new antibacterial with potent activity against Gram-positive bacteria but thought to have limited potency against Gram-negative bacteria.

**Objectives:** This study evaluated the antibacterial activity of MGB-BP-3 under nutrient-limited, physiologically relevant conditions to better mimic *in vivo* environments.

**Methods:** Minimum inhibitory concentrations (MICs) were determined against a panel of ESKAPEE pathogens in both rich (Cation Adjusted Mueller-Hinton Broth, CA-MHB) and nutrient-limited media (Roswell Park Memorial Institute, RPMI-1640 + 10% Luria-Bertani, LB). Synergy with efflux pump inhibitors and membrane permeabilizers was evaluated via checkerboard assays. Permeability was assessed using a fluorescent DNA-binding cell permeant (Hoechst 33342) accumulation assay. MICs were determined in the presence of different concentrations of ions, using both wild-type and efflux-compromised strains. *In vivo* efficacy was assessed using the *Galleria mellonella* infection model.

**Results:** The activity of MGB-BP-3 against Gram-negative bacteria was improved by shifting the medium from rich to nutrient-limited conditions. Specifically, it showed potent activity against E. coli in nutrient-limited media (MIC 3.1 µM) but was inactive in rich media (MIC 700 µM). Synergy with efflux inhibitors occurred mainly in rich media, indicating barriers to intracellular accumulation were not present in nutrient-limited conditions. Increased permeability correlated with enhanced susceptibility and efflux-deficient strains were more susceptible, confirming the role of efflux pumps in resistance. MGB-BP-3 provided dose-dependent protection in *G. mellonella* larvae infected with *E. coli* or *S. aureus*. For example, at concentrations of 12.5 µM and 100 µM, MGB-BP-3 was able to protect the larvae by more than 70% over 5 days of treatment, respectively.

**Conclusions:** Nutrient-limited media better reveal MGB-BP-3’s activity against Gram-negative bacteria and align with *in vivo* efficacy, highlighting the need for host-mimicking conditions in antibiotic screening.

## 1. Introduction

Finding new antibiotics is crucial for combating antimicrobial resistance (AMR) (1). Traditionally, the process of finding antibiotics has used nutrient-rich media, such as cation adjusted Mueller-Hinton broth (CA-MHB). The rapid bacterial growth supported by these media facilitate the screening process of new antibiotics (2). However, these conventional methods often fail to mimic the more nutrient-limited *in vivo* environment, which may lead to inaccurate predictions between *in vitro* screening and *in vivo* efficacy (3).

Several recent *in vitro* studies using nutrient-limited media have discovered active compounds that were inactive in commonly used traditional rich media. Luna *et al*. have shown that rifabutin has better antibacterial activity against highly resistant *Acinetobacter baumannii*, in nutrient-limited media, with an MIC 200-fold lower compared to rich media and markedly improved the survival rate of infected mice (4). In another study, Buyck *et al*. have shown that *Pseudomonas aeruginosa* is more susceptible to macrolide antibiotics in nutrient-limited media, compared to rich media, due to decreased expression of efflux pumps and increased outer-membrane permeability (5).

MGB-BP-3 is a novel antibacterial compound of the minor groove binders’ group that has shown potent activity against Gram-positive bacteria and has successfully completed a Phase IIa clinical trial for the treatment of *Clostridioides difficile (6)*. MGB-BP-3 is thought to be ineffective against Gram-negative bacteria due to its poor intracellular accumulation, which is a result of efficient efflux systems and/or poor membrane permeation, as shown earlier in nutrient rich media, CA-MHB (6).

Given that nutrient-limited conditions are thought to alter the permeability of the bacterial envelope, we were interested to investigate whether nutrient-limited conditions would result in an improvement of activity for MGB-BP-3 against Gram-negative bacteria.

In this study, we evaluated the antibacterial activity of MGB-BP-3 against the ESKAPEE pathogens (7,8) under different growth conditions, from nutrient-defined to nutrient-rich media. Moreover, we explored the protective effect of MGB-BP-3 against *E. coli* infection using the *Galleria mellonella* model, a more physiologically relevant, nutrient-limited environment.

This study highlights the importance of physiologically relevant in vitro screening methods and simple in vivo models for more accurate prediction of antimicrobial efficacy.

## 2. Materials and Methods

### 2.1. Test compounds and other materials

MGB-BP-3 was synthesized as previously described (9). Polymyxin B Nonapeptide (PMBN, P2076) and Phenyl-arginine-beta-naphthylamide (PaβN, P4157) were purchased from Sigma-Aldrich. All other materials that were used in this study were purchased from Fisher Scientific: dehydrated Mueller-Hinton Broth (MHB, Catalogue number CM0405B), Select Agar (Catalogue number 30391023), Luria-Bertani Broth (LB, Catalogue number 12780029), Roswell Park Memorial Institute Medium (RPMI-1640, Catalogue number 61870010), Hoechst 33342 (Catalogue number 62249) and DMSO (Catalogue number 85190). All materials were handled and stored according to the manufacturer’s instructions.

### 2.2. Bacterial Strains, Growth Conditions and *Galleria mellonella* larvae

Bacterial strains of *Staphylococcus aureus ATCC 43300, Enterococcus faecalis ATCC 51299, Klebsiella pneumoniae* ATCC 700603, *E. coli* ATCC 25922, *P. aeruginosa* ATCC 27893 and *Acinetobacter baumannii* ATCC 19606 were obtained from the American Type Culture Collection (ATCC). Clinical isolates of *E. coli 170020100, E. coli 170020013, E. coli 150310983, E. coli 150164245 (6)* were obtained from NHS Lanarkshire. Laboratory model strains of *E. coli MG1655, E. coli ΔacrB and E. coli D408A* were kindly provided by Dr. Jessica Blair from University of Birmingham. Initially, a single bacterial colony from a freshly streaked MHA or LB agar plate was transferred into CA-MHB or LB and incubated overnight at 35±2°C with shaking at 140 rpm. The next day, the overnight culture was diluted 1:100 in CA-MHB, LB, RPMI, or RPMI supplemented with 10% of LB medium, and incubated at 37°C with shaking at 140 rpm until the culture reached exponential growth phase, with an optical density at 600 nm (OD_600_) of 0.5. Optical density was measured by TECAN Spark™ multimode (Tecan Austria GmbH) or HIDEX Sense (Hidex Oy) microplate readers.

Wax moth larvae (*Galleria mellonella*) were purchased from Livefood UK Ltd (Rooks Bridge, UK) for *in vivo* studies.

### 2.3. Minimum Inhibitory Concentration

The Minimum Inhibitory Concentrations (MICs) were determined by broth microdilution in 96-well microplates as described by CLSI (18). Initially, 100 µL of the respective medium containing 5 × 10^5^ CFU/ml bacteria (OD_600_=0.5) were dispensed into the wells of flat-bottom 96-well plates in columns 1-11. Columns 11 and 12 served as positive and negative controls, containing only bacteria with medium and medium without bacteria, respectively. In column 1, 198 µL of bacterial suspension and 2 µL of MGB-BP-3 solution (10 mM in DMSO) were added. In column 2, 198 µL of the respective culture medium and 2 µL of drug working concentration were added, followed by two-fold serial dilutions of the MGB-BP-3 across columns 2-10 (100 µM-0.2 µM). The OD_600_ of the plates was read initially and after incubation at 35±2°C in a static incubator for 20 hours. The MIC was defined as the lowest concentration of the compound that inhibited bacterial growth. While OD_600_ readings were taken and corroborated with OD_600_ measurements by subtracting OD_600t=0h_ from OD_600t=20h_. Each experiment was performed in triplicate. To determine the effect of specific divalent cations on antibacterial activity, the medium was supplemented individually with a range of concentrations of calcium, magnesium, or iron. Sterile stock solutions of calcium chloride (CaCl2), magnesium sulfate (MgSO4), and iron (II) sulfate (FeSO4) were prepared and added to the basal medium prior to bacterial inoculation. MICs were then determined for each cation-supplemented condition using the standard broth microdilution protocol.

### 2.4. Fractional Inhibitory Concentration Index

The synergy between MGB-BP-3 and other compounds, including PMBN and PAβN, was evaluated by a checkerboard broth microdilution method (10) using either CA-MHB or RPMI+10% LB.

Serial two-fold dilutions of MGB-BP-3 and other compounds were prepared in either CA-MHB or RPMI+10% LB from their stock solutions. For synergy testing, 25 µL of MGB-BP-3 dilutions were added to the wells containing PMBN or PABN. Then 50 µL of double-concentrated bacterial stock (2 × 10^5^ CFU/mL) was added to each well, bringing the final volume to 100 µL. The bacterial inoculum was prepared as per the MIC protocol. The dilutions were arranged in a flat bottom 96-well microtiter plate with each well containing a unique combination of the two drugs at different concentrations (8). OD_600_ was measured before and after the plates were incubated in a static incubator at 35±2°C for 18±2 hours and the fractional inhibitory concentration index (FICI) was calculated by the following formula:

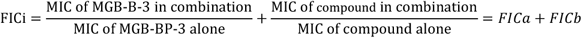

FICI values of ≤ 0.5, 0.5 < FICI ≤ 1, 1.0 < FICI ≤ 4, and FICI > 4.0 indicate synergy, additive effects, indifference, and antagonism, respectively (11). In cases where the MICs exceeded the highest concentration tested, the highest concentration in the dilution series was used to determine the FICI, and the result reported as ≤ the calculated value.

### 2.5. Bactericidal Activity Assay

The time-kill assay was conducted to evaluate whether MGB-BP-3 has bactericidal or bacteriostatic activity in RPMI medium supplemented with 10% LB. An *E. coli* 25922 culture was prepared from an exponential phase growth and adjusted to a final concentration of 1 × 10^6^ CFU/mL in 15 mL Falcon tubes containing 5 mL of RPMI+10%LB. The cultures were then exposed to various multiples of the MGB-BP-3 MICs (0.5, 1, 5, 10). Culture tubes were incubated at 35±2°C in a shaking incubator and aliquots of 100 mL were taken at 0, 1, 3, 5, 7, 20 and 24 hours post-inoculation. These were 10-fold serially diluted in sterile 0.9% saline solution and plated on MH agar to establish the bacterial concentration as colony forming units per mL (CFU/mL) according to the drop plate method (12, 13). Plates were incubated at 35±2°C for 24 hours. Colony-forming units (CFU) were counted at each time point and the results plotted as log_10_ CFU/mL versus time. A reduction of ≥3 log_10_ CFU/mL compared to the initial inoculum is considered bactericidal, while bacteriostatic activity is below this threshold.

### 2.6. Assessment of Intracellular Accumulation

*E. coli, P. aeruginosa, K. pneumoniae, A. baumannii* strains were cultured overnight at 37°C in CA-MHB and RPMI+10%LB separately, then the next morning they were used to inoculate fresh medium and incubated for an additional 3-5 hours to bring the culture back to the exponential growth phase. Bacteria were then harvested by centrifugation at 4000 × *g*, resuspended in PBS, and adjusted to an OD_600_ of 0.2. Aliquots of 180 µL were transferred into individual columns (n=4 replicates per strain) of a flat-bottomed black 96-well plate. Column 1 served as the PBS blank; columns 2-5 contained heat-inactivated strains (used as controls, with bacterial heat-inactivated in a water bath at 90°C for 10 min after adjustment to OD_600_ = 0.2); and columns 6-9 were allocated for test strains. Hoechst 33342 dye (20 µL of 50 µM) was added to each well to achieve a final concentration of 5 µM. Fluorescence was measured from the top of the plate with 10 flashes per well (TECAN Spark™ multimode plate reader; excitation: 355 nm; emission: 460 nm). Data were processed in Excel: mean fluorescence values were calculated after blank subtraction. Fluorescence intensity is presented as a percentage of the signal from heat-inactivated strain, reflecting membrane permeability. Experiments were performed in triplicate. Statistical significance was assessed using a paired Student’s t-test to compare the outer-membrane permeability of each bacterial strain when cultured in RPMI+10% LB versus CA-MHB. The same experiment was also carried out for efflux pump compromised *E. coli* strains (*E. coli* ΔacrB and *E. coli* D408A) *(14,15)*.

### 2.7. *Galleria mellonella* Larvae Infection Model

Ethical approval for this work was received from the University of Glasgow School of Biodiversity, One Health & Veterinary Medicine Research Ethics Committee (ref EA04/24). Larvae at their final instar stage (the last larval stage before pupation), weighing between 250-350 mg, were kept at room temperature in darkness and used within one week of receipt. Bacterial cultures (*E. coli* ATCC 25922 and *S. aureus* ATCC 43300) were incubated overnight in LB broth and sub-cultured the next day to reach an OD_600_ of 0.5. The cultures were then centrifuged at 5000 × *g* for 10 minutes and the pellets were resuspended in sterile PBS to achieve the desired inoculum concentration of 5 × 10^6^ CFU/mL and 5 × 10^8^ CFU/mL concentration for *E. coli* and *S. aureus*, respectively. Each larva (10 per group) was injected with 10 µL of the bacterial suspension (approximately 5 × 10^4^ CFU/larva for *E. coli* and 5 ×10^6^ CFU/larva for *S. aureus*) into the hind left proleg by using a needle (26S gauge, 50.8 mm length, point style 4 (30º), Hamilton, Timis, Romania) and a syringe (1750 RN 500 mL, Hamilton). Two hours post-infection, the larvae were treated with 10 µL of a single concentration of MGB-BP-3 injected into the hind right proleg. The larvae were then incubated at 37°C in a 90-mm plastic Petri dish and monitored for survival rate over a 5-day period. Larvae were considered dead when they did not move in response to stimulus with a pipette tip (16, 17). Survival data were recorded and analysed using Kaplan-Meier survival method (18), with statistical significance determined using the log-rank test (p < 0.05) (19). Those larvae that started to pupate during the monitoring process were excluded from the analysis. For generating the MGB-BP-3 toxicity, *G. mellonella* larvae (10 larvae per group) only received a range of doses of the compound and monitored for survival for five days.

### 2.8. Statistical analysis

All statistics were performed using GraphPad Prism 9.5.1 version.

## 3. Results and Discussion

Previous studies have shown that MGB-BP-3 has significant antibacterial activity against Gram-positive bacteria, with low MIC values (< 1 µM). However, no measurable activity against Gram-negative bacteria, including *E. coli*, due to poor intracellular accumulation was identified (6). As previous work had only been carried out using nutrient-rich media, specifically CA-MHB, we hypothesised that MGB-BP-3 might exhibit antibacterial effects against *E. coli* under physiologically relevant conditions.

To test this hypothesis, we used RPMI medium supplemented with 10% LB, a combination that has been suggested to better mimic physiological conditions relevant to human use (20). Similar to a study by Kumaraswamy *et al* (21), we found that RPMI alone did not provide a sufficient fast growth rate and density of the bacteria to enable us to carry out antimicrobial screening against ESKAPEE pathogens (Figure S1). However, supplemented with 10% of LB we achieved growth rates that were relatively comparable to those in CA-MHB (Figure S1).

We next determined the MIC of MGB-BP-3 against a panel of ESKAPE pathogens and *E. coli* in LB broth, MHB, CA-MHB and RPMI+10%LB (Table 1). As expected, MGB-BP-3 had low MICs against both Gram-positive organisms in all media, and no measurable activity against Gram-negative bacteria in the nutrient-rich media (MHB, CA-MHB and LB broth). However, MGB-BP-3 exhibited significant antibacterial activity in RPMI+10%LB, with MIC values of 3.1 µM against *E. coli* and 50 µM against *A. baumannii*. These results indicate that the culture medium composition can influence compound potency against these bacterial strains, particularly *E. coli*. At certain points, the MIC of MGB-BP-3 against Gram-negative bacteria was determined to be >100 µM due to the compound’s solubility limit, and we were unable to assess any potential improvements beyond this concentration. The promising MIC against *E. coli* ATCC 25922 prompted us to assess MGB-BP-3 against a small panel of multi-drug-resistant clinical isolates (6). For two of these (170020100 and 170020013), in RPMI-10%LB, MGB-BP-3 also had comparable MICs, while it didn’t show any measurable activity against the remaining two isolates (150310983 and 150164245).

**Table 1.** MIC of MGB-BP-3 (µM) against ESKAPEE pathogens at different culture conditions. *E. coli* clinical isolates described in ref (6)

| Organism | Strain | RPMI+10%LB | MHB | CA-MHB | LB Broth |
| --- | --- | --- | --- | --- | --- |
| *S. aureus* | ATCC 43300 | 0.19 | 0.31 | 0.31 | 0.31 |
| *E. faecalis* | ATCC 51299 | 0.31 | 0.39 | 0.39 | 0.39 |
| *A. baumannii* | ATCC 19606 | 50 | >100 | >100 | >100 |
| *K. pneumoniae* | ATCC 700603 | >100 | >100 | >100 | >100 |
| *P. aeruginosa* | ATCC 27893 | >100 | >100 | >100 | >100 |
| *E. coli* | ATCC 25922 | 3.1 | >100 | >100* | >100 |
| *E. coli* | 170020100 | 3.1 | >100 | >100 | >100 |
| *E. coli* | 150310983 | >100 | >100 | >100 | >100 |
| *E. coli* | 150164245 | >100 | >100 | >100 | >100 |
| *E. coli* | 170020013 | 12.5 | >100 | >100 | >100 |

- In one case, we tested *E. coli* ATCC 25922 with a higher concentration of MGB-BP-3, and the MIC was approximately 700 μM.

We suspected that the lower activity of MGB-BP-3 in rich media against many Gram-negative bacteria was due to lower intracellular accumulation, and so we conducted synergy experiments, combining MGB-BP-3 with PAβN and PMBN (Table 2, Figure S5). PMBN is a well-known membrane permeabilizing agent (24), whereas PAβN has a dual mechanism, primarily acting as an efflux pump inhibitor and also enhancing membrane permeability at high concentrations (25). The combination of MGB-BP-3 with either PAβN or PMBN showed better synergistic activity in CA-MHB compared to RPMI+10%LB. For example, in CA-MHB, the FICI values for all tested bacteria were lower than 0.5 except for *K. pneumoniae*, indicating strong synergy. In contrast, in RPMI+10%LB, the FICIs were greater, or equal, compared to CA-MHB, showing reduced synergy. The lower synergy in nutrient-limited media suggests a lower permeability barrier for MGB-BP-3.

**Table 2.** Fractional Inhibitory Concentration indices (FICIs) of MGB-BP-3 in combination with PABN and PMBN against various standard Gram-negative ESKAPE bacterial strains in CA-MHB and RPMI+10%LB media. FICI<0.5 indicates synergy.

| Medium | Compound | *E. coli* | *K. pneumoniae* | *A. baumannii* | *P. aeruginosa* |
| --- | --- | --- | --- | --- | --- |
| CA-MHB | PAβN | 0.06 | 0.09 | 0.09 | 0.26 |
|  | PMBN | 0.09 | 2 | 0.08 | 0.2 |
| RPMI+10%LB | PAβN | 1.06 | 2 | 0.15 | 1.5 |
|  | PMBN | 0.5 | 2 | 0.18 | 1 |

To further probe the origin of the lower MICs observed in RPMI+10%LB, we investigated the hypothesis that variable cation concentrations between the test media (RPMI+10%LB and CA-MHB) were responsible. We studied the impact of varying concentrations of calcium (Ca^2+^, magnesium (Mg^2+^), and iron (Fe^2+^) on the antibacterial activity of MGB-BP-3 against *E. coli* by supplementing basal RPMI+10%LB with each cation individually (Figure. 1).

**Figure 1.**
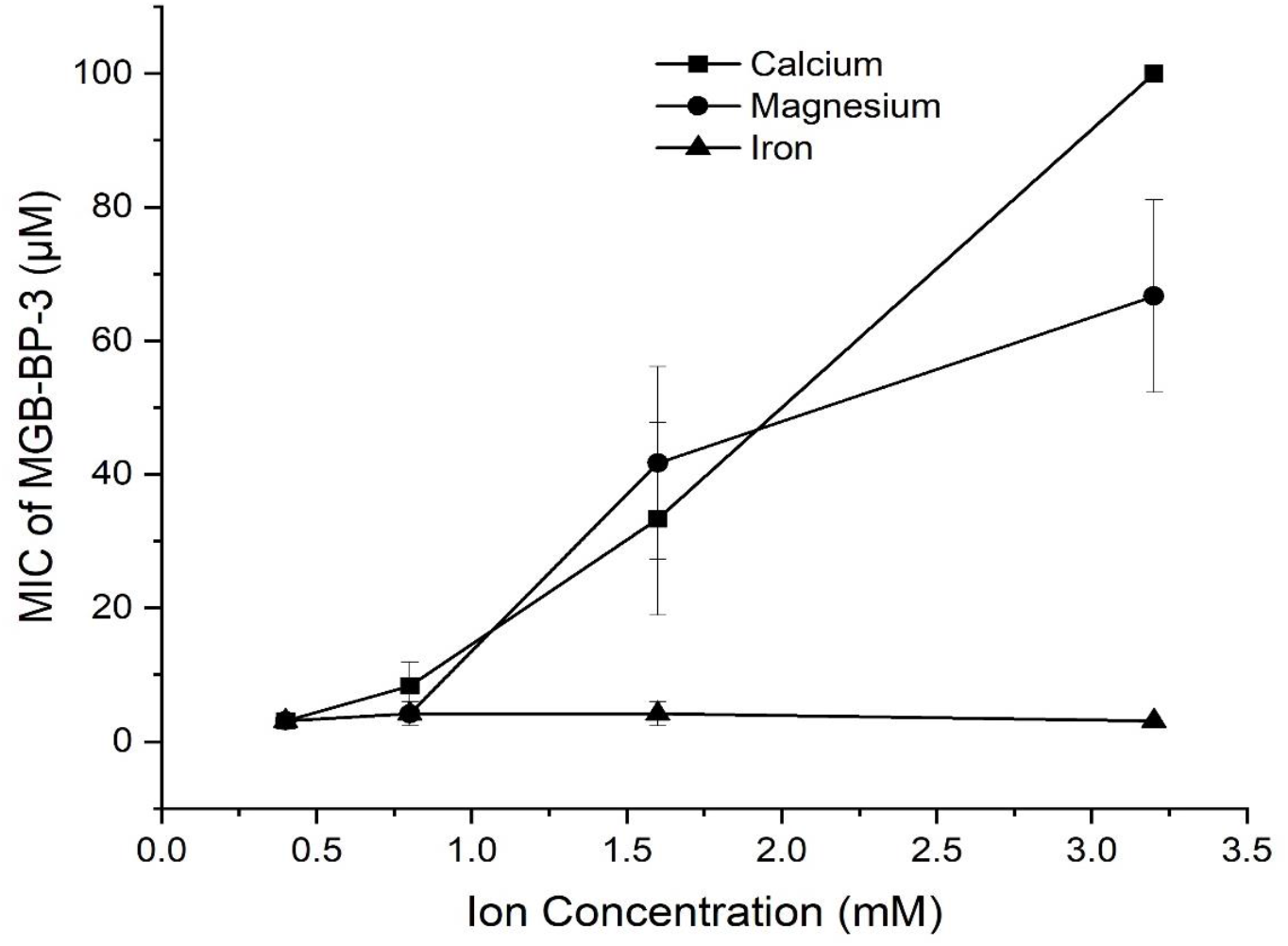
Effect of calcium (Ca^2+^), magnesium (Mg^2+^) and ferrous (Fe^2+^) concentrations on the antibacterial activity of MGB-BP-3 against *E. coli* ATCC 25922 in RPMI+10%LB. Increasing cation concentrations to 0.8 mM, 1.6 mM, and 3.2 mM resulted in higher MIC values, indicating reduced antibacterial activity of MGB-BP-3.

**Figure 2.**
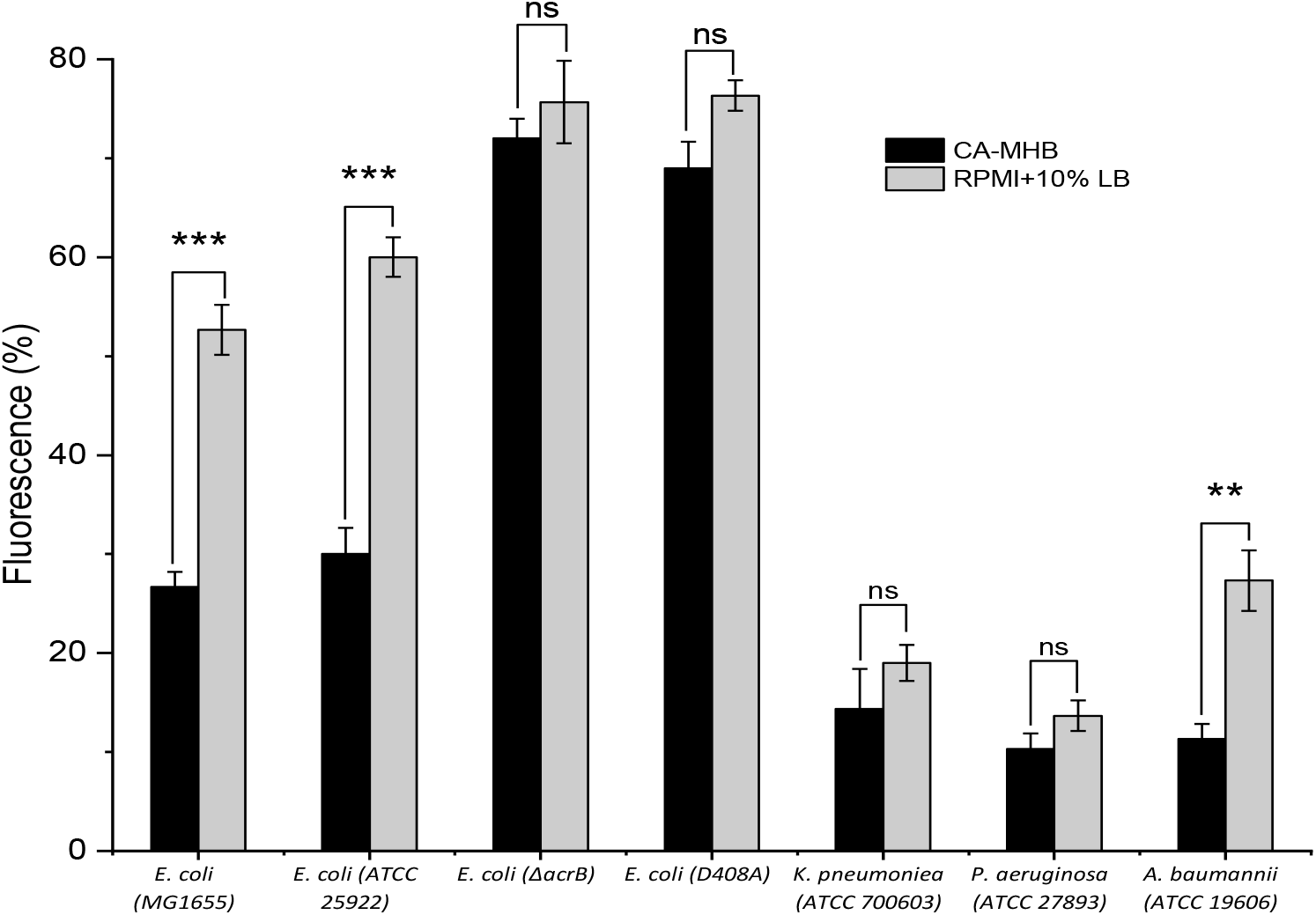
Relative accumulation of Hoechst 33342 in Gram-negative bacterial strains cultured in RPMI+10%LB and CA-MHB media. Fluorescence intensity is presented as a percentage of the signal from heat-inactivated strain, reflecting membrane permeability. Data represent the mean fluorescence from three independent experiments, each comprising four technical replicates per strain (*E. coli-1: E. coli MG1655, E. coli-2: E. coli ATCC25922, E. coli-3 ΔacrB, E. coli-4: E. coli D408A, K. pneumoniae ATCC 700603, P. aeruginosa ATCC 27893 and Acinetobacter baumannii ATCC 19606)*. All data are presented as the mean (±SD) from three independent replicates (n=3). A two-sample independent t-test was performed for each strain to compare the two media. Significance is indicated as ** (p < 0.01) or *** (p < 0.001) and non-significant is indicated as (ns).

The results showed that increasing cation concentrations (Ca^2+^ and Mg^2+^) significantly increased the MIC of MGB-BP-3. However, increasing the concentration of iron in the media did not affect the MIC. The increased MIC of MGB-BP-3 in the presence of calcium and magnesium ions is likely due to their role in stabilizing the bacterial outer membrane (26), thereby reducing its permeability to MGB-BP-3. Iron ions were not observed to affect membrane permeability in the same way and thus did not alter the MIC under these experimental conditions.

To understand the contribution of the effects of efflux on MGB-BP-3 activity, we determined MICs against wild type and efflux pump compromised *E. coli* MG1655 strains (Table 3). The AcrAB-TolC efflux pump system plays a crucial role in mediating multidrug resistance in various bacteria, including Enterobacteriaceae. Over 33-fold reduction in MGB-BP-3’s MIC was observed against mutant strains with compromised efflux, either via a gene knock-out (Δ*acr*B) or deactivation due to a non-synonymous mutation (D408A) in the inner membrane transporter AcrB when tested in CA-MHB medium, and a 7-fold reduction in RPMI+10%LB medium. This suggests that efflux mediated through AcrAB-TolC is a component of reduced efficacy of MGB-BP-3 in *E. coli* in nutrient-rich media. Although, the difference in fold reduction (>33 vs 7-fold) between the wild type and efflux compromised strains could suggest that the nutrient-limited media increases the permeability of the *E. coli* membrane.

**Table 3.** MICs of MGB-BP-3 against wild type and efflux pump compromised (ΔacrB and D408A) mutants of *E. coli* MG1655 in CA-MHB and RPMI+10%LB media.

| Compound ID | MIC (μM) in CA-MHB | | | MIC (μM) in RPMI+10%LB | | |
| --- | --- | --- | --- | --- | --- | --- |
| | *E. coli* MG1655 | *E. coli* ΔacrB | *E. coli* D408A | *E. coli* MG1655 | *E. coli* ΔacrB | *E. coli* D408A |
| **MGB-BP-3** | >100 | 3.1 | 3.1 | 3.1 | 0.4 | 0.4 |

We next investigated membrane differences in RPMI+10%LB medium compared to CA-MHB using a Hoechst 33342 accumulation assay (Figure 3). It should be noted that Hoechst 33342 is known to be an efflux substrate, and so this assay cannot deconvolute permeability from efflux (23, 27). The increase in fluorescence intensity was statistically significantly greater in the nutrient-limited media for both wild type *E. coli* and *A. baumannii* strains, but not for *P. aeruginosa, K. pneumoniae* or the efflux-compromised *E. coli* strains. *P. aeruginosa* and *K. pneumoniae* are known to possess highly efficient efflux systems and have reduced porin expression, factors which contribute to their lower intracellular compound accumulation (22, 24). These species-specific traits appear to override the influence of media composition, as *E. coli* and *A. baumannii* responded with increased accumulation of Hoechst 33342 when cultured in RPMI+10%LB. The high accumulation observed in efflux pump-compromised *E. coli* strains under both media conditions was likely due to Hoechst 33342 being an efflux substrate (23) resulting in its expulsion from the wild-type cells. These permeability and efflux profiles are in accordance with the MIC values of MGB-BP-3 (Table 1 and 3) against these strains in both media and support a strong correlation between the membrane’s barrier function and antimicrobial susceptibility.

**Figure 3.**
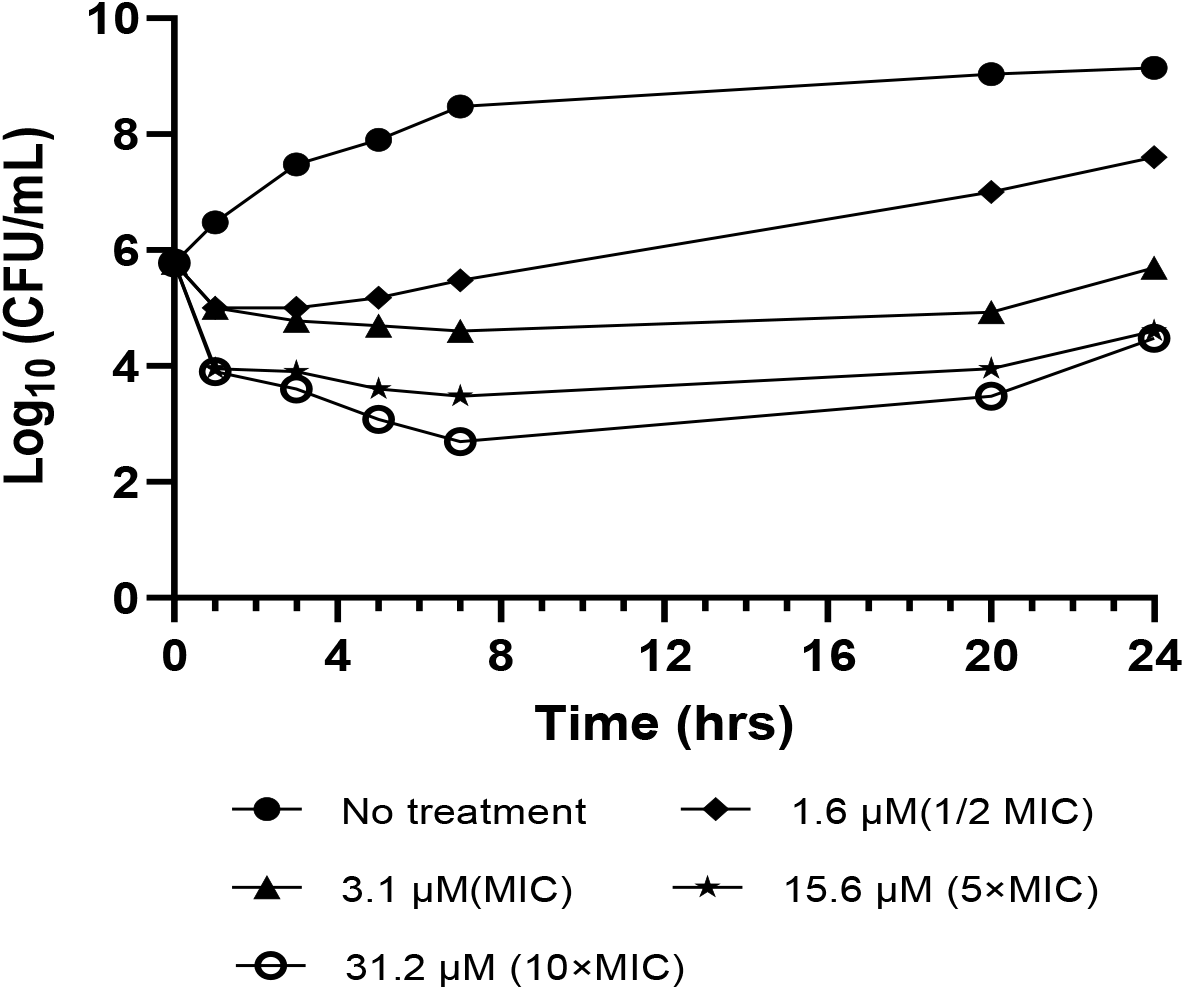
Viable counts for cultures of *E. coli* 25922. Results are shown as No treatment control and cultures treated at 35±2° C for 24 hours, with MGB-BP-3 concentration corresponding to 0.5, 1, 5 and 10 times the MIC value determined.

Given that MGB-BP-3 was effective against several stains of *E. coli*, we decided to probe its mechanism further. We performed a time-kill assay to assess its bactericidal potential. *E. coli* was exposed to MGB-BP-3 concentrations ranging from 0.5 to 10 times the determined MIC values over an incubation period of 24 hours (Figure 3).

Bacterial killing, i.e. bactericidal activity of MGB-BP-3 was concentration-dependent as shown on time-kill curves (Figure 3). Incubation of *E. coli* cells with an MGB-BP-3 concentration of 1/2MIC initially resulted in a gradual decrease in viable bacterial cell count. However, after a short time (∼4 h), CFU counts began to increase, and after 24 hours of incubation growth at the bottom of the tube and biofilm formation were observed (Figure 3). For f MGB-BP-3 at or over the MIC concentrations (≥3.1 µM), bacterial suspensions reverted to growth and CFU counts began to increase by 20 hours, and the reduction in bacterial cell counts was less than 3 log_10_ CFU/mL after 24 hours of incubation, indicating a bacteriostatic effect at this time point. Bactericidal effect was observed at 31.2 mM MGB-BP-3 after 7 hours incubation (Figure 3).

The bactericidal action of 10x MIC or greater concentrations has been observed even for commonly considered bactericidal antimicrobials such as b-lactams, fluoroquinolones and aminoglycosides, especially in the treatment of Gram-positive infections (28). There are other factors, such as increasing heterogeneity in bacterial populations beyond exponential growth phase, and with this associated drug penetration limits (planktonic vs biofilm cultures) or kinetics of target inactivation (e.g. reduced DNA replication in stationary growth phase) that require further exploration (31).

Given that we observed different MICs for MGB-BP-3 against *E. coli*, but not *S. aureus*, depending on whether rich media or nutrient-limited media were used, we were interested to investigate the compound’s *in vivo* efficacy. Therefore, we conducted a *Galleria mellonella* larvae infection experiment (Figure 4).

**Figure 4.**
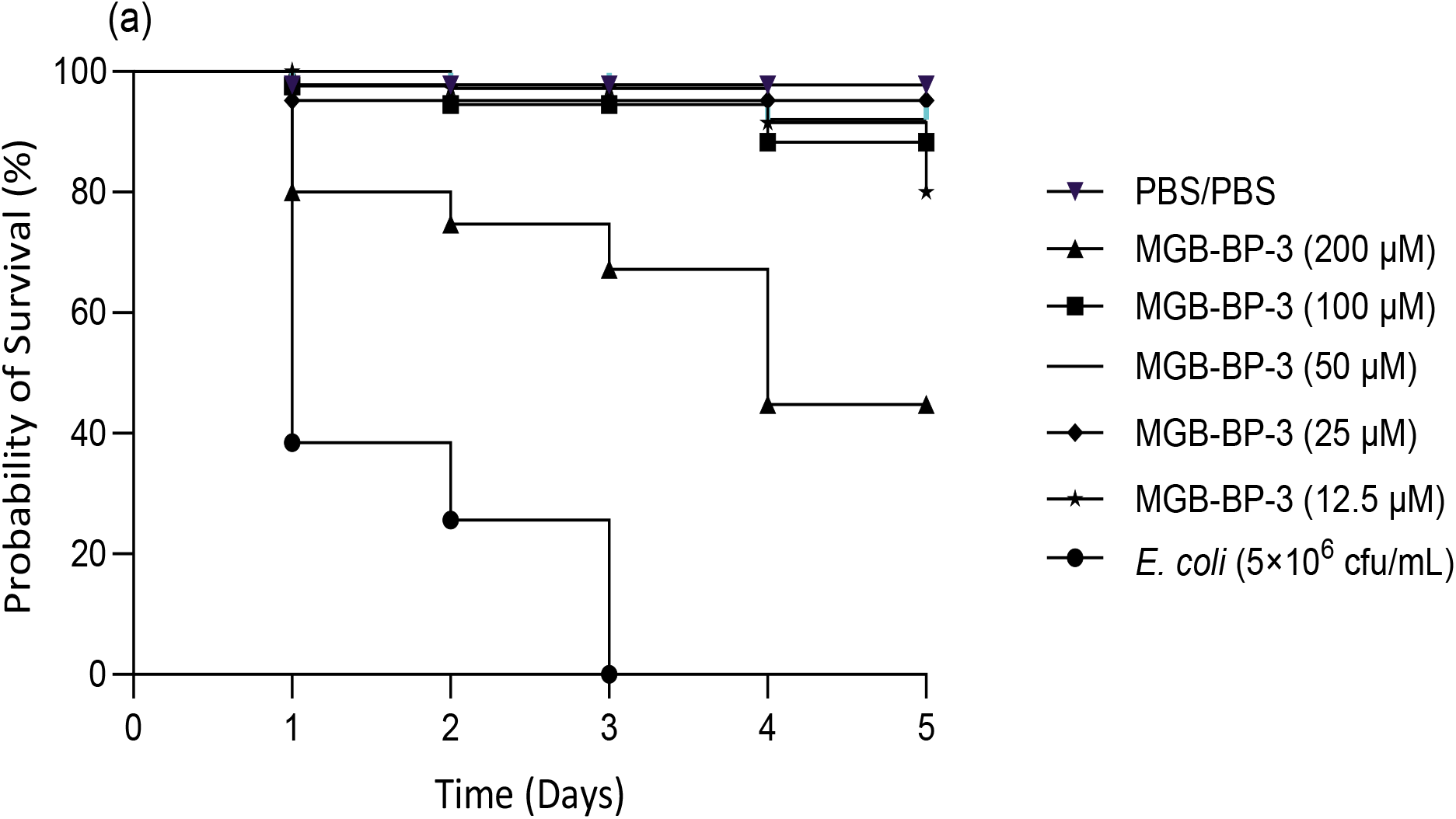
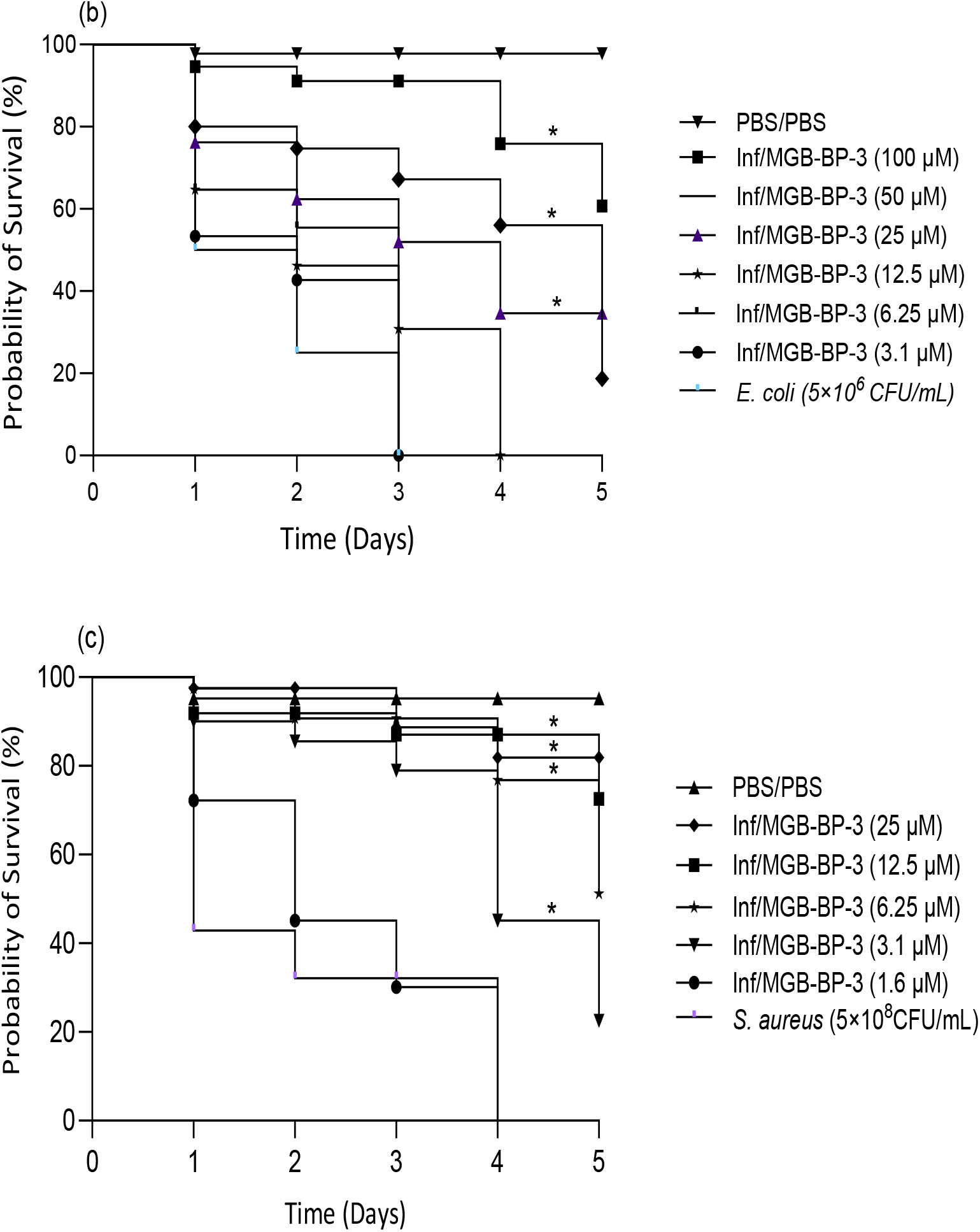
(a) Toxicity effects of MGB-BP-3 on G. mellonella larvae at various concentrations, monitored over 5 days. (b) Survival probability of G. mellonella larvae after 5 days of treatment with MGB-BP-3 following infection with 5×10^6^ CFU/mL of E. coli ATCC 25922 strain. (c) Survival probability of G. mellonella larvae after 5 days of treatment with MGB-BP-3 following infection with 5×10^8^ CFU/mL of S. aureus. Larvae were administered either PBS (control-treated) or MGB-BP-3 at various doses and incubated at 37°C. A single dose of the MGB-BP-3 treatment was given 2 hours post-infection. The number of surviving larvae was recorded every day for a total of 5 days. The uninfected PBS/PBS group represents larvae that were sham injected with sterile PBS (to serve as a control for the physical injection) and subsequently treated with sterile PBS. Statistical analyses were conducted using the log-rank test, with a significance level set at (p < 0.05) for multiple comparisons. All experiments were performed in triplicate, with a sample size of (n = 30)

Initially, we assessed the toxicity of MGB-BP-3 against *G. mellonella* larvae. At the highest dose, 50 mg/kg (64×MIC or 200 µM), half of the larvae died after 4 days, indicating some toxicity effect at this dose. However, at a dose of 25 mg/kg (32×MIC or 100 µM) no significant toxic effects were observed (Figure 4a). To compare the *in vivo* protective effects of MGB-BP-3 with its *in vitro* activities in both rich and nutrient-limited media, *E. coli* and *S. aureus* were chosen for infection in *G. mellonella*. The lethal dose of each bacterium was determined to be 5×10^6^ CFU/mL for *E. coli* after 3 days (Figures S2 and S3) and 5×10^8^ CFU/mL for *S. aureus* after four days (Figures S2 and S4).

Following the initial experiments, the larvae were infected with a single lethal dose of each bacterium and two hours post-infection, treated with MGB-BP-3 at concentrations ranging from 25 mg/kg (100 µM) to 0.8 mg/kg (3.1 µM) for *E. coli* and 6.2 mg/kg (25 µM) to 0.8 mg/kg (0.3 µM) for *S. aureus* infection. The larvae were monitored for 5 days. The results demonstrated a dose-dependent protective effect of MGB-BP-3 against both bacterial strains. For larvae infected with *E. coli*, a concentration of 25 µM of MGB-BP-3 protected approximately 40% of the larvae for 5 days. A concentration of 12.5 mg/kg (50 µM) provided 60% protection after 4 days. Approximately 70% of the larvae remained alive after 5 days of treatment with 25 mg/kg (100 µM) of MGB-BP-3 compared to a control group – infected but treated with PBS, which showed significantly lower survival rates, and all larvae dying after three days of infection (Figure 4b). However, MGB-BP-3 protected the larvae infected with *S. aureus* at lower concentrations, starting at 0.8 mg/kg (3.1 µM) which provided around 30% protection after 5 days of infection and increasing the MGB-BP-3 dose enhanced the protection effect as at the concentration of 3.1mg/kg (12.5 mM) and 6.2 mg/kg (25 µM )about 90% of the infected larvae remained alive after 5 days of treatment with either dose (Figure 4c). The in vivo dose required to achieve 60% larval survival translated to a remarkably similar multiple of the in vitro MIC for both pathogens. For *S. aureus*, the protective dose of 6.25 µM was 31-fold its MIC (0.2 µM) in MHB-2, while for *E. coli*, the protective dose of 100 µM was approximately 32-fold its MIC in RPMI (3.1 µM).

These findings indicate that MGB-BP-3 exhibits significant protective effects in a dose-dependent manner against both *E. coli* and *S. aureus* infections in the *G. mellonella* model. Interestingly, the MICs obtained in the nutrient-limited media were more in line with the data obtained in the *G. mellonella* infection model. Although this model has been established for *in vivo* infections and antimicrobial treatment testing (30, 31) antimicrobial *in vivo* efficacy and the measured MIC values are affected by various other factors, yet to be studied for MGB-BP-3.

## 4. Conclusion

Several studies have shown that bacterial growth in host-mimicking conditions can reveal the potential of antibiotics that were previously thought to be ineffective against certain strains. For instance, rifabutin exhibited high potency against extensively drug-resistant *A. baumannii* in nutrient-limited media, unlike in standard, rich media (4). In another study, azithromycin showed improved efficacy against multidrug-resistant *P. aeruginosa* in RPMI-1640 compared to traditional media. Culture medium influences bacterial gene expression, membrane permeability, and resistance/tolerance mechanisms which all impact antibiotic susceptibility testing (5). Therefore, host-mimicking media enhance the accuracy of antimicrobial susceptibility testing, especially for intracellular-targeting antibiotics (4, 5).

Our research demonstrates that MGB-BP-3 has a significant antibacterial activity against *E. coli* in RPMI+10% LB medium, with a MIC of 3.1 µM versus over 100 µM in rich media (Table 1). This suggests that medium composition affects MGB-BP-3’s efficacy, particularly against Gram-negative bacteria. Synergy experiments with PAβN and PMBN indicate that permeability barriers and efflux pumps reduce MGB-BP-3’s effectiveness in rich media (Table 2), while varying cation levels (calcium and magnesium) increase MIC of MGB-BP-3 by stabilizing the bacterial outer membrane in nutrient-limited media (Figure 1). Moreover, screening MGB-BP-3 against wild type and efflux pump-compromised *E. coli* MG1655 strains revealed that the AcrAB-TolC efflux pump system significantly contributes to the reduced efficacy of MGB-BP-3 in rich media, with a >33-fold reduction in MIC in MHB medium compared to a 7-fold reduction in RPMI medium (Table 3).

These *in vitro* experiments provide evidence that the high susceptibility of *E. coli* to MGB-BP-3 in nutrient-limited media, compared to broths, is likely due to increased accumulation of the compound within the bacteria. This is likely attributed to changes in the outer membrane integrity caused by the medium in contact with the bacterial cells, along with a reduction in their active efflux pumps as we observed in the permeability assay and Hoechst 33342 uptake in RPMI+10% LB compared to CA-MHB (Figure 2). These findings align with prior research on rifabutin, azithromycin, and other antibiotics emphasizing the value of host-mimicking conditions for assessing antibiotics with intracellular targets, especially against Gram-negative bacteria (4,5).

The *in vitro* MIC data obtained in the nutrient-limited RPMI+10%LB, but not CA-MHB, are more congruent with the *G. mellonella* infection model data for both *E. coli* and *S. aureus*. Ultimately, this study contributes to the emerging narrative that MIC testing in nutrient-rich media may not be suitable to accurately predict *in vivo* outcomes, which is particularly important when considering new antibiotic drug discovery.

Within this context, we also reveal that MGB-BP-3, a compound previously thought to have no clinically relevant activity against Gram-negative bacteria, has therapeutically relevant MIC against *E. coli* using nutrient-limited media and shows promise in clearing an *E. coli* infection in the *G. mellonella* larvae model. Consequently, MGB-BP-3 may be worth considering for further investigation in a murine model of infection.

## 5. Author Contributions

KE designed and carried out all experiments and contributed to manuscript preparation. LCY, KO, JMAB and FJS carried out executive manuscript preparation, management and research conception.

## 6. Acknowledgements

For the purpose of open access, the author(s) has applied a Creative Commons Attribution (CC BY) licence to any Author Accepted Manuscript version arising from this submission.

## 7. Conflicts of Interest

The authors declare the following competing financial interest(s): FJS is part of revenue sharing agreements with their University relating to the Strathclyde minor groove binder project. Additionally, FJS has financial interests through shares in the company, Rostra Therapeutics. All other authors: none to declare.

## References

1. Tang, K. W. K., Millar, B. C., & Moore, J. E. (2023). Antimicrobial resistance (AMR). British Journal of Biomedical Science, 80, 11387. 10.3389/bjbs.2023.11387.

2. Wiegand, I., Hilpert, K., & Hancock, R. (2008). Agar and broth dilution methods to determine the minimal inhibitory concentration (MIC) of antimicrobial substances. Nature Protocols, 3(2), 163–175 10.1038/nprot.2007.521.

3. Belanger, C. R., & Hancock, R. E. W. (2021). Testing physiologically relevant conditions in minimal inhibitory concentration assays. Nature Protocols, 16(8), 3761–3774. 10.1038/s41596-021-00572-8.

4. Luna, B., Trebosc, V., Lee, B., et al. (2020). A nutrient-limited screen unmasks rifabutin hyperactivity for extensively drug-resistant Acinetobacter baumannii. Nature Microbiology, 5(9), 1134–1143. 10.1038/s41564-020-0737-6.

5. Buyck, J. M., Plésiat, P., Traore, H., Vanderbist, F., Tulkens, P. M., & Van Bambeke, F. (2012). Increased susceptibility of Pseudomonas aeruginosa to macrolides and ketolides in eukaryotic cell culture media and biological fluids due to decreased expression of oprM and increased outer-membrane permeability. Clinical Infectious Diseases, 55(4), 534–542. 10.1093/cid/cis473.

6. Hind, C., et al. (2022). Insights into the spectrum of activity and mechanism of action of MGB-BP-3. ACS Infectious Diseases, 8(12), 2552–2563. 10.1021/acsinfecdis.2c00445.

7. Louis B Rice, Federal Funding for the Study of Antimicrobial Resistance in Nosocomial Pathogens: No ESKAPE, The Journal of Infectious Diseases, Volume 197, Issue 8, 15 April 2008, Pages 1079–1081. 10.1086/533452.

8. Partridge SR, Kwong SM, Firth N, Jensen SO. Mobile Genetic Elements Associated with Antimicrobial Resistance. Clin Microbiol Rev. 2018 Aug 1;31(4): e00088–17. doi: 10.1128/CMR.00088-17.

9. Brooke, D. P., McGee, L. M. C., Giordani, F., Cross, J. M., Khalaf, A. I., Irving, C., et al. (2021). Truncated S-MGBs: Towards a parasite-specific and low aggregation chemotype. RSC Medicinal Chemistry, 12(9), 1391–1401. 10.1039/D1MD00110H.

10. Bellio, P., Fagnani, L., Nazzicone, L., & Celenza, G. (2021). New and simplified method for drug combination studies by checkerboard assay. MethodsX, 8, 101543. 10.1016/j.mex.2021.101543.

11. Odds, F. C. (2003). Synergy, antagonism, and what the chequerboard puts between them. Journal of Antimicrobial Chemotherapy, 52(1), 1. 10.1093/jac/dkg301.

12. National Committee for Clinical Laboratory Standards. (1999). Methods for determining bactericidal activity of antimicrobial agents; approved guidelines, M26-A (Vol. 19). NCCLS.

13. Silva, F., Lourenço, O., Queiroz, J. A., & Domingues, F. C. (2011). Bacteriostatic versus bactericidal activity of ciprofloxacin in Escherichia coli assessed by flow cytometry using a novel far-red dye. Journal of Antibiotics (Tokyo), 64(4), 321–325. 10.1038/ja.2011.5.

14. Coldham, N. G., Webber, M., Woodward, M. J., & Piddock, L. J. V. (2010). A 96-well plate fluorescence assay for assessment of cellular permeability and active efflux in Salmonella enterica serovar Typhimurium and Escherichia coli. Journal of Antimicrobial Chemotherapy, 65, 1655–1663. 10.1093/jac/dkq169.

15. Richmond, G. E., Chua, K. L., & Piddock, L. J. V. (2013). Efflux in Acinetobacter baumannii can be determined by measuring accumulation of H33342 (bis-benzamide). Journal of Antimicrobial Chemotherapy, 68, 1594–1600. 10.1093/jac/dkt052.

16. Hill, L., Veli, N., & Coote, P. J. (2014). Evaluation of Galleria mellonella larvae for measuring the efficacy and pharmacokinetics of antibiotic therapies against Pseudomonas aeruginosa infection. International Journal of Antimicrobial Agents, 43(3), 254–261. 10.1016/j.ijantimicag.2013.11.001.

17. Fletcher, M., McCormack, A., Parcell, B. J., & Coote, P. J. (2023). Combination therapy with ciprofloxacin and pentamidine against multidrug-resistant Pseudomonas aeruginosa: Assessment of in vitro and in vivo efficacy and the role of resistance–nodulation–division (RND) efflux pumps. Antibiotics, 12(8), 1236. 10.3390/antibiotics12081236.

18. Bland, J. M., & Altman, D. G. (1998). Survival probabilities (the Kaplan-Meier method). BMJ, 317(7172), 1572. 10.1136/bmj.317.7172.1572.

19. Bland, J. M., & Altman, D. G. (2004). The logrank test. BMJ, 328(7447), 1073. 10.1136/bmj.328.7447.1073.

20. Belanger, C. R., Lee, A. H., Pletzer, D., Dhillon, B. K., Falsafi, R., & Hancock, R. E. W. (2020). Identification of novel targets of azithromycin activity against Pseudomonas aeruginosa grown in physiologically relevant media. Proceedings of the National Academy of Sciences of the United States of America, 117(52), 33519–33529. 10.1073/pnas.2007626117.

21. Kumaraswamy, M., Lin, L., Olson, J., Sun, C. F., Nonejuie, P., Corriden, R., Döhrmann, S., Ali, S. R., Amaro, D.,Rohde, M., Pogliano, J., Sakoulas, G., & Nizet, V. (2016). Standard susceptibility testing overlooks potent azithromycin activity and cationic peptide synergy against MDR Stenotrophomonas maltophilia. Journal of Antimicrobial Chemotherapy, 71(5), 1264–1269. 10.1093/jac/dkv487.

22. Leus, I. V., & Zgurskaya, H. I. (2025). No two are alike: On the role of Klebsiella pneumoniae permeability barriers in antibiotic susceptibility and persistence. Antimicrobial Agents and Chemotherapy, 69, e00085–25. 10.1128/aac.00085-25.

23. Whittle, E. E., Orababa, O., Osgerby, A., Siasat, P., Element, S. J., Blair, J. M. A., & Overton, T. W. (2024). Efflux pumps mediate changes to fundamental bacterial physiology via membrane potential. mBio, 15, e02370–24. 10.1128/mbio.02370-24.

24. Haim Tsubery, Itzhak Ofek, Sofia Cohen, and Mati Fridkin (2000). Structure−Function Studies of Polymyxin B Nonapeptide: Implications to Sensitization of Gram-Negative Bacteria. Journal of Medicinal Chemistry 2000 43 (16), 3085–3092. 10.1021/jm0000057.

25. Lamers RP, Cavallari JF, Burrows LL (2013) The Efflux Inhibitor Phenylalanine-Arginine Beta-Naphthylamide (PAβN) Permeabilizes the Outer Membrane of Gram-Negative Bacteria. PLOS ONE 8(3):e60666. 10.1371/journal.pone.0060666.

26. Luke A. Clifton, Maximilian W. A. Skoda, Anton P. Le Brun, Filip Ciesielski, Ivan Kuzmenko, Stephen A. Holt, and Jeremy H. Lakey. (2015). Effect of Divalent Cation Removal on the Structure of Gram-Negative Bacterial Outer Membrane Models. Langmuir 31 (1), 404–412. 10.1021/la504407v.

27. Richmond GE, Chua KL, Piddock LJ. Efflux in Acinetobacter baumannii can be determined by measuring accumulation of H33342 (bis-benzamide). J Antimicrob Chemother. 2013 Jul;68(7):1594–600 10.1093/jac/dkt052.

28. G. A. Pankey, L. D. Sabath, Clinical Relevance of Bacteriostatic versus Bactericidal Mechanisms of Action in the Treatment of Gram-Positive Bacterial Infections, Clinical Infectious Diseases, Volume 38, Issue 6, 15 March 2004, Pages 864–870, 10.1086/381972.

29. Brauner, A., Fridman, O., Gefen, O. et al. Distinguishing between resistance, tolerance and persistence to antibiotic treatment. Nat Rev Microbiol 14, 320–330 (2016). 10.1038/nrmicro.2016.34.

30. Andrew P. Desbois, Peter J. Coote, Wax moth larva (Galleria mellonella): an in vivo model for assessing the efficacy of antistaphylococcal agents, Journal of Antimicrobial Chemotherapy, Volume 66, Issue 8, August 2011, Pages 1785–1790, 10.1093/jac/dkr198.

31. Dougal H. Adamson, Vasare Krikstopaityte, Peter J. Coote, Enhanced efficacy of putative efflux pump inhibitor/antibiotic combination treatments versus MDR strains of Pseudomonas aeruginosa in a Galleria mellonella in vivo infection model, Journal of Antimicrobial Chemotherapy, Volume 70, Issue 8, August 2015, Pages 2271–2278, 10.1093/jac/dkv111.

