## Supplementary Data for "Nutrient-limited conditions reveal the activity of a minor groove binder, MGB-BP-3, against *Escherichia coli*"

### Contents

**1. Growth rates in different media (Page 2)**

**2. Checkerboard assays (Page 2)**

**3. Calibration of OD600 to CFU/mL, and lethality of infection dose (Page 9)**

### 1. Growth rates in different media

A single colony of *E. coli ATCC*25922 and *S. aureus* ATCC43300 inoculated into 10 mL of CA-MHB were grown overnight to stationary phase (18 hrs) at 37°C in a shaking incubator with 140 rpm. The next day bacteria were washed twice with PBS and re-suspended in tubes containing 25 mL of CA-MHB, RPMI+10%LB or RPMI to an initial $OD_{600}$ of 0.05. Tubes were then placed in a shaking incubator with 141rpm at 37°C and re-growth measured by spectrophotometer by measuring $OD_{600}$ at selected time for up to 24h (**Figure S1**).

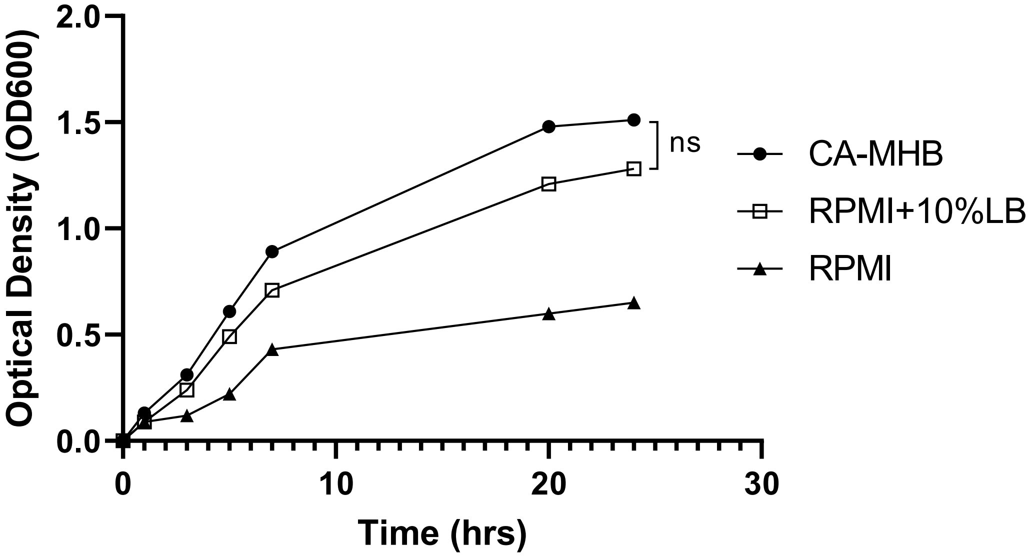


**Figure S1.** *E. coli* growth was monitored in three different media: CA-MHB (rich medium), RPMI, and RPMI supplemented with 10% LB. Bacterial growth was measured by turbidity using a spectrophotometer at OD600 at the following time points: 1, 3, 5, 7, 20, and 24 hours. Statistical analysis using independent samples t-tests revealed no significant difference (ns) in growth between CA-MHB and RPMI supplemented with 10% LB.

### 2. Checkerboard assays

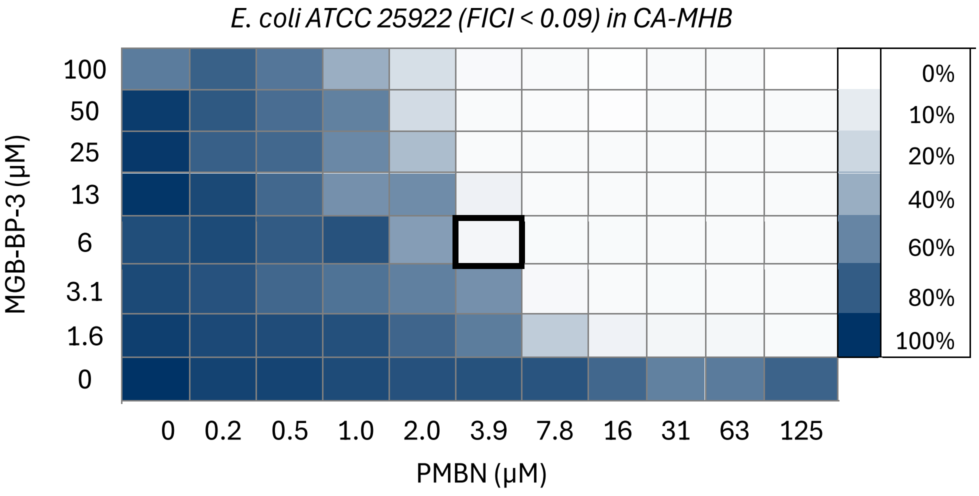


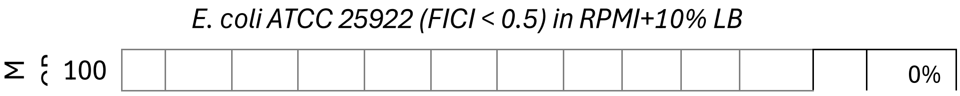

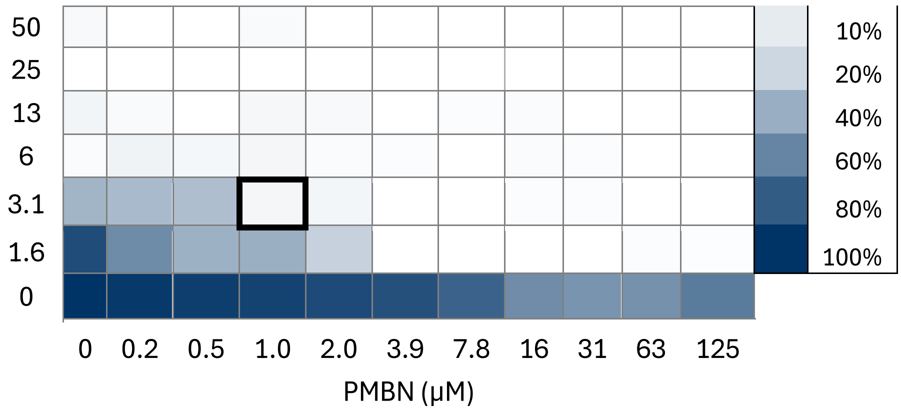


*E. coli ATCC 25922 (FICI <1.03) in RPMI+ 10% LB*

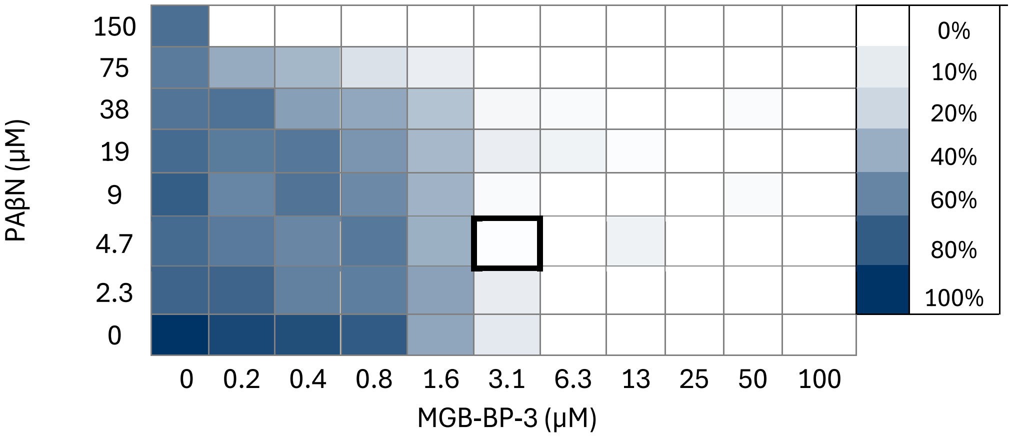


*E. coli ATCC 25922 (FICI <0.09) in CA-MHB*

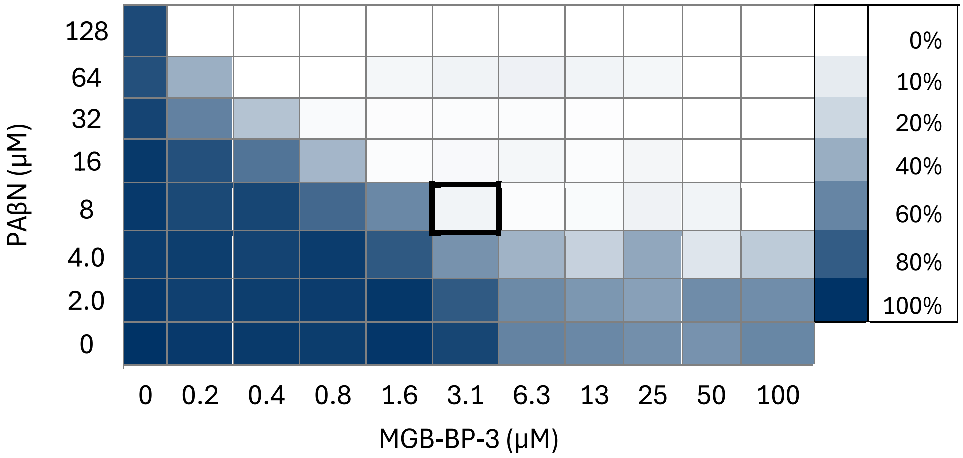


*K. pneumoniae ATCC 700603 (FICI < 2) in RPMI+10%LB*

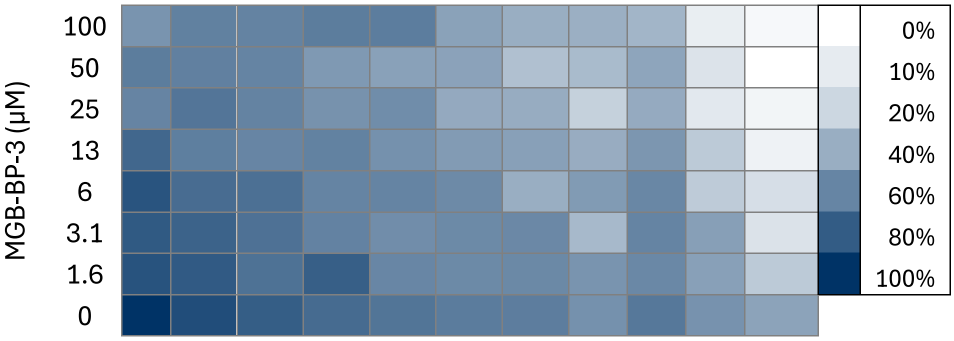

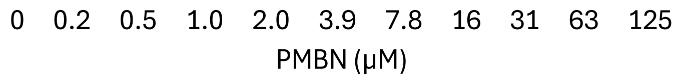


*K. pneumoniae ATCC 700603 (FICI < 2) in CA-MHB*

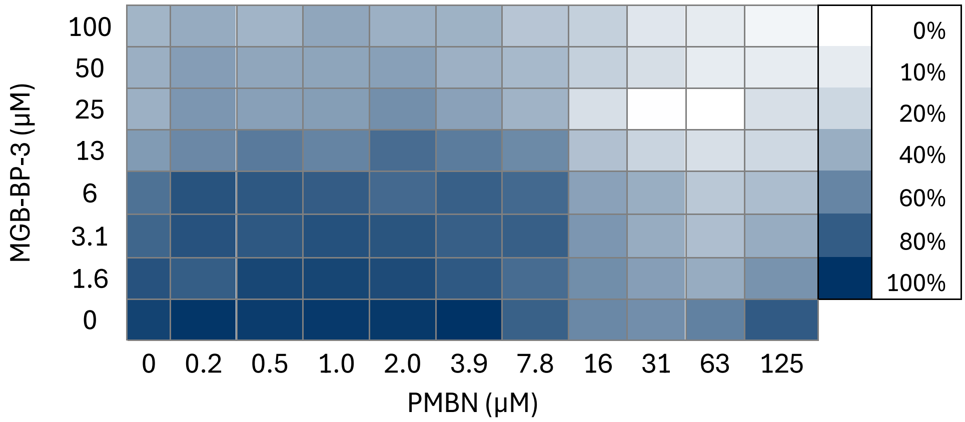


*K. pneumoniae ATCC 700603 (FICI <2) in RPMI+10%LB*

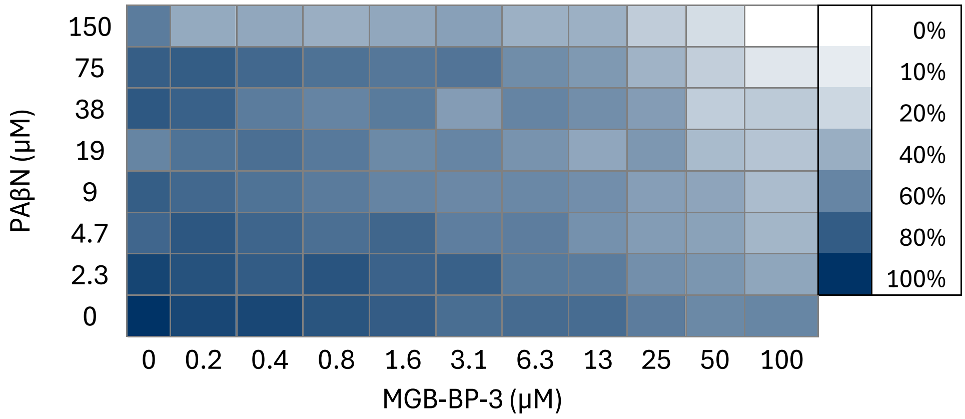


*K. pneumoniae ATCC 700603 (FICI <0.09) in CA-MHB*

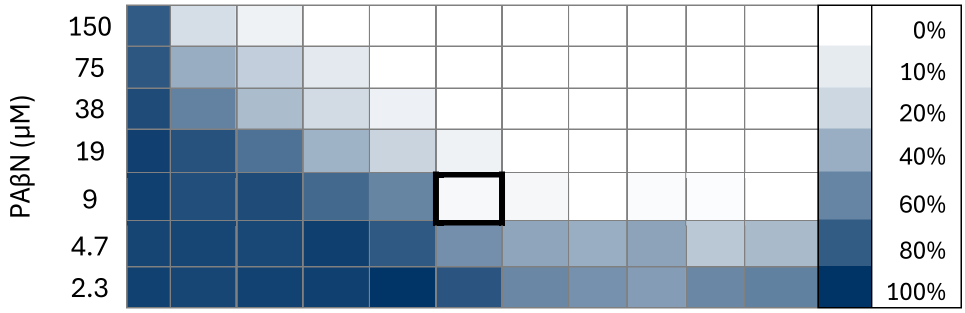

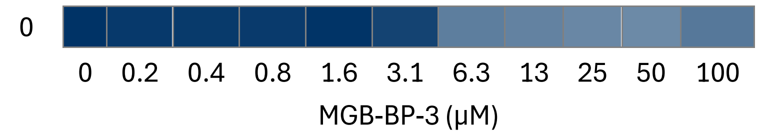
0
0 0.2 0.4 0.8 1.6 3.1 6.3 13 25 50 100
MGB-BP-3 (µM)

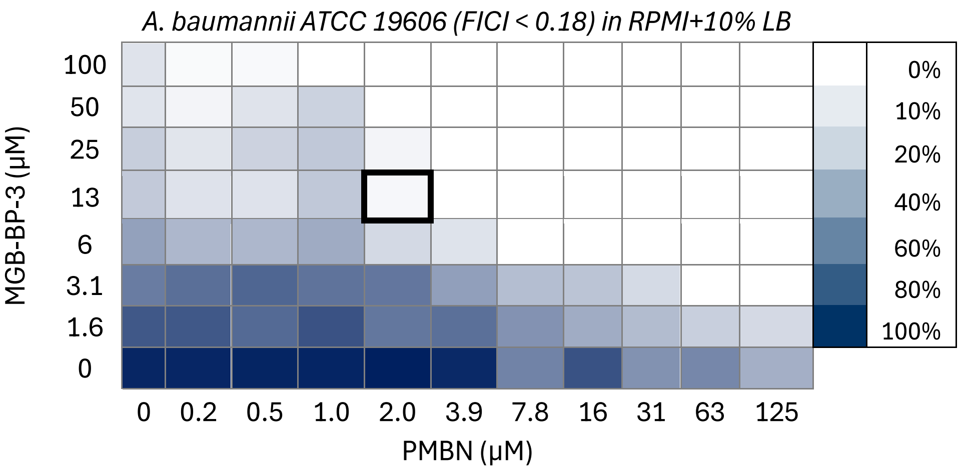
A. baumannii ATCC 19606 (FICI < 0.18) in RPMI+10% LB
MGB-BP-3 (µM)
100 50 25 13 6 3.1 1.6 0
0 0.2 0.5 1.0 2.0 3.9 7.8 16 31 63 125
PMBN (µM)
0% 10% 20% 40% 60% 80% 100%

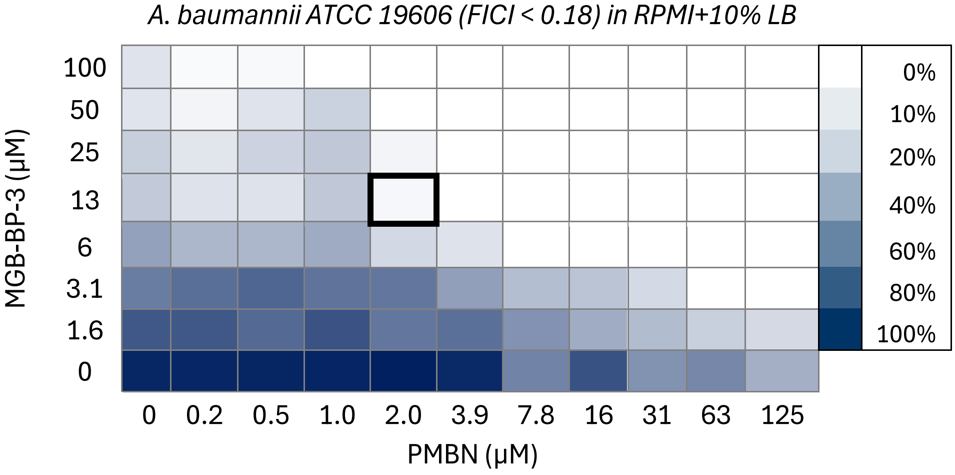
A. baumannii ATCC 19606 (FICI < 0.18) in RPMI+10% LB
MGB-BP-3 (µM)
100 50 25 13 6 3.1 1.6 0
0 0.2 0.5 1.0 2.0 3.9 7.8 16 31 63 125
PMBN (µM)
0% 10% 20% 40% 60% 80% 100%

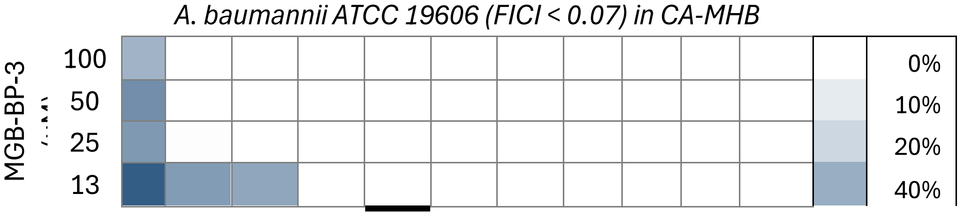
A. baumannii ATCC 19606 (FICI < 0.07) in CA-MHB
MGB-BP-3
100 50 25 13
0% 10% 20% 40%

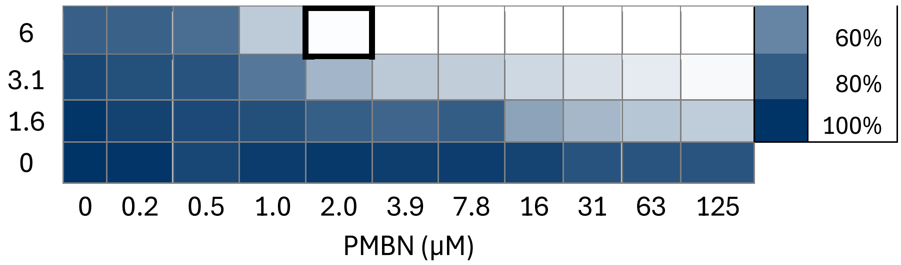


*A. baumannii ATCC 19606 (FICI <0.06) in CA-MHB*

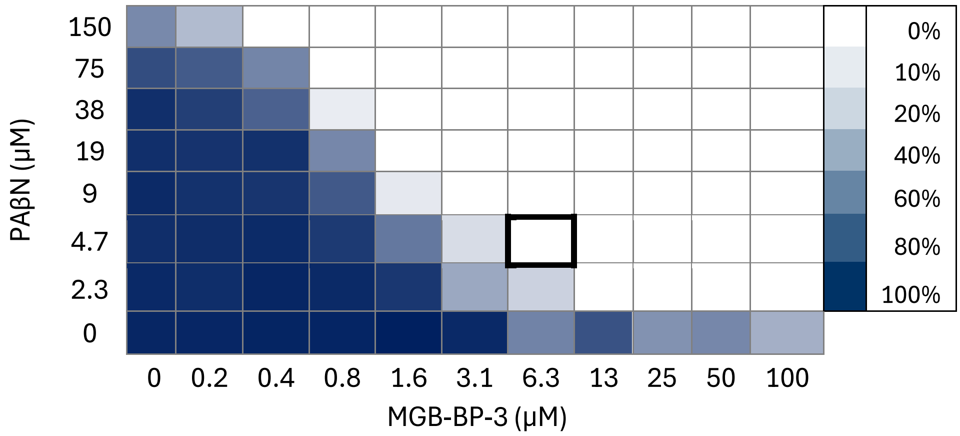


*K. pneumoniae ATCC 700603 (FICI < 2) in RPMI+10%LB*

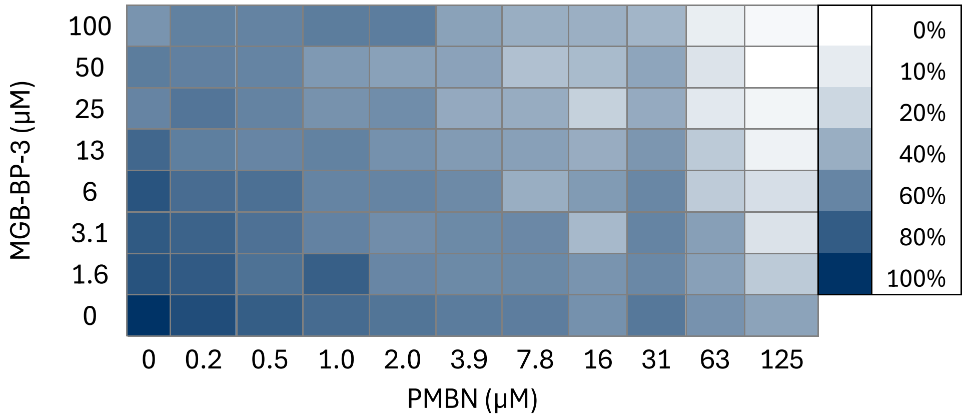


*A. baumannii ATCC 19606 (FICI <0.15) in RPMI+10%LB*

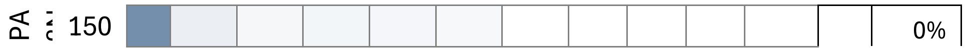

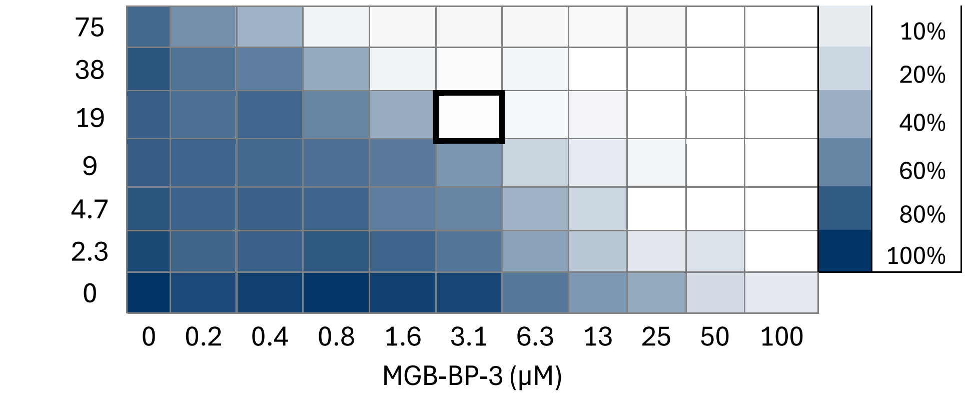


*A. baumannii ATCC 19606 (FICI <0.06) in CA-MHB*

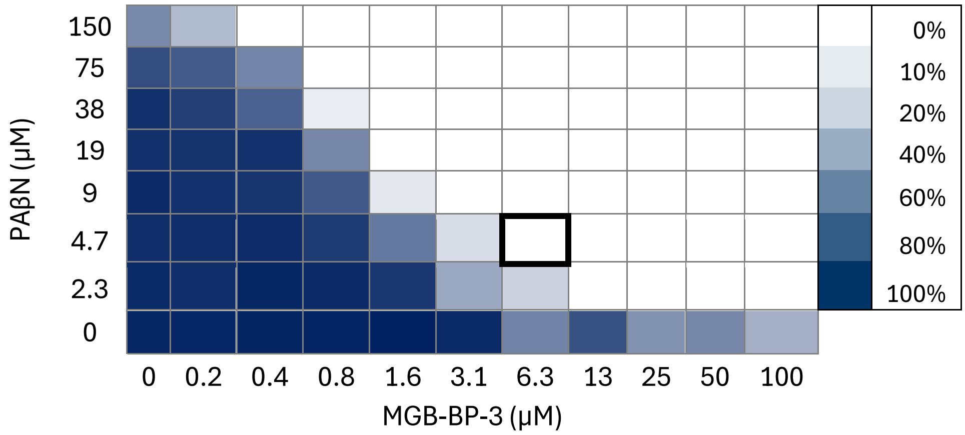


*A. baumannii ATCC 19606 (FICI < 0.07) in CA-MHB*

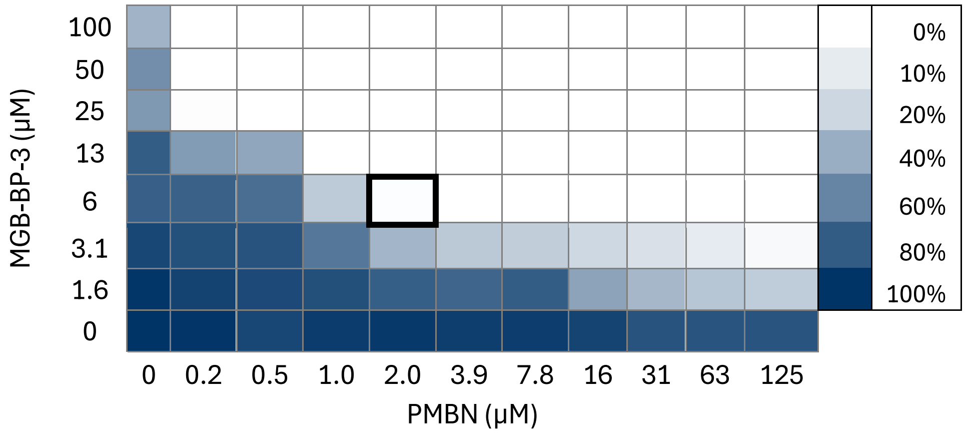


*A. baumannii ATCC 19606 (FICI < 0.18) in RPMI+10% LB*

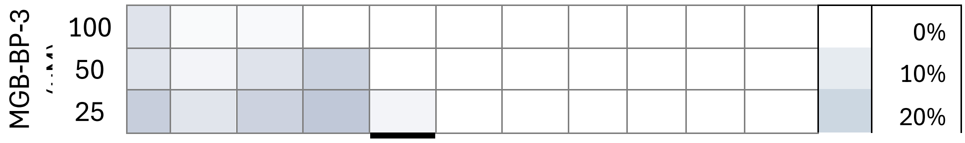

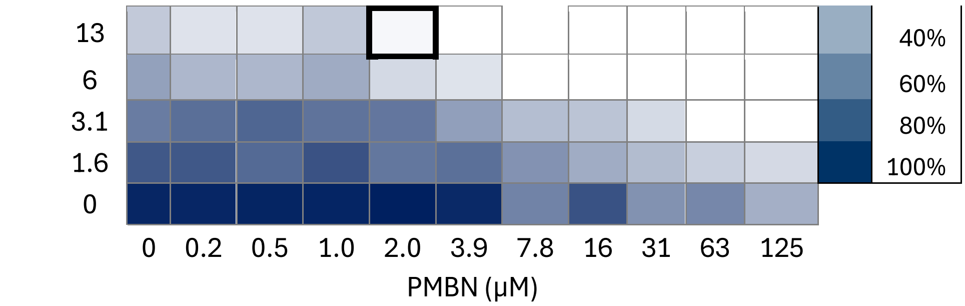


*P. aeruginosa ATCC 27893 (FICI <1) in RPMI+10%LB*

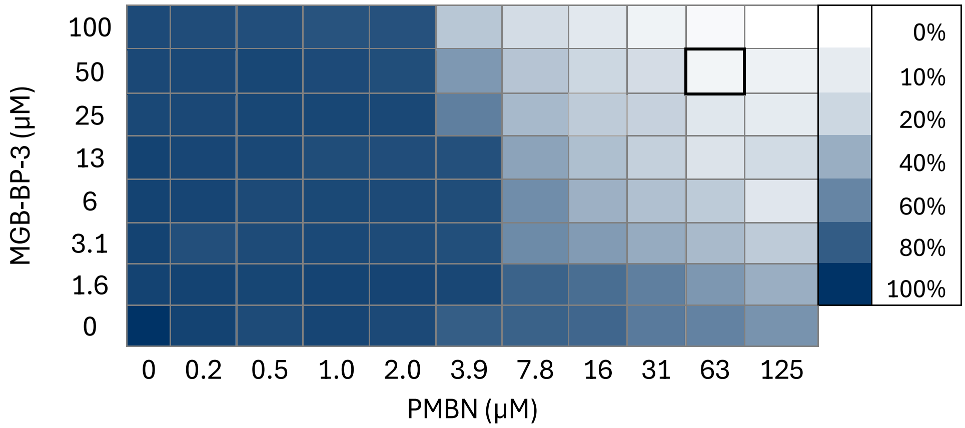


*P. aeruginosa ATCC 27893 (FICI < 0.2) in CA-MHB*

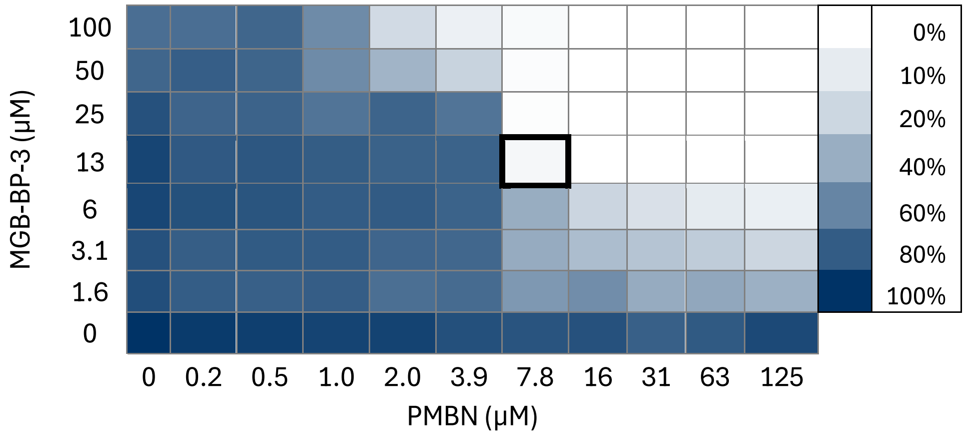


*P. aeruginosa ATCC 27893 (FICI <1.8) in RPMI+10%LB*

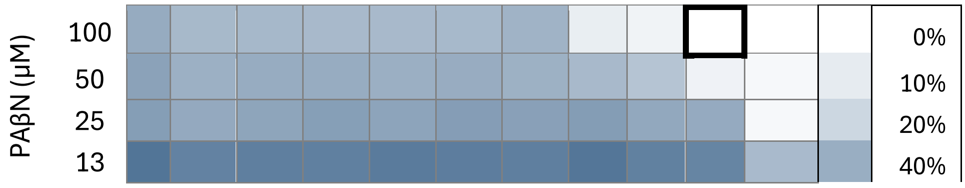

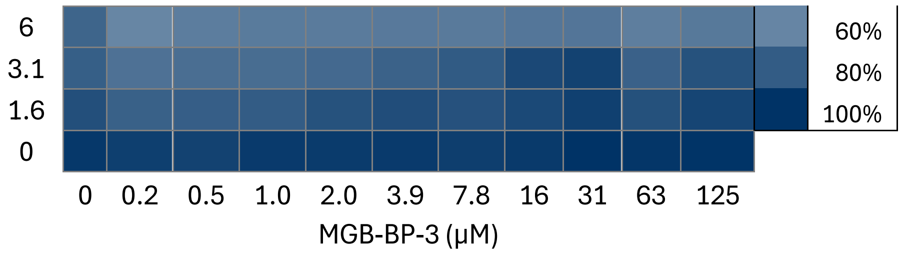


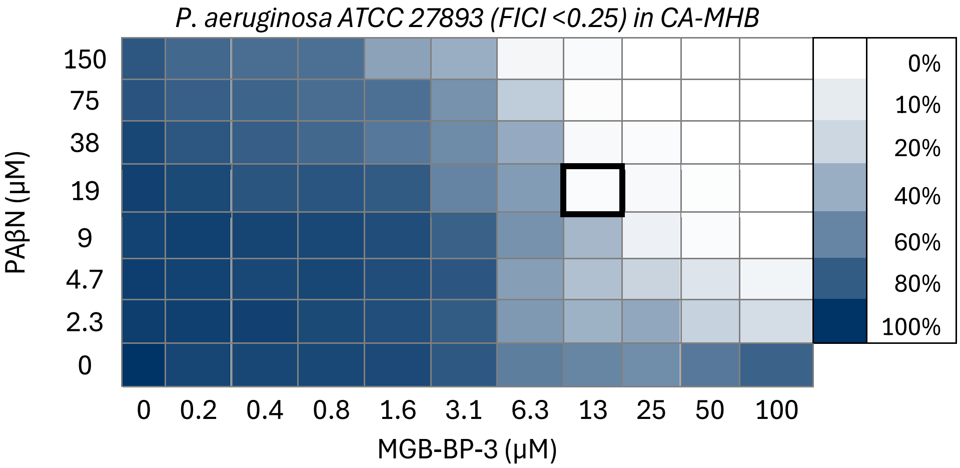


**Figure S5.** Checkerboard assays of MGB-BP-3 in combination with PaβN or PMBN against Gram-negative bacteria from ESKAPE pathogen panel in different media condition. Fractional inhibitory concentration indices (FICIs) from checkerboard assays indicate significant synergy for values <0.5. Calculated FICIs for all strains are mentioned above the panels.

#### 3. Calibration of OD600 to CFU/mL, and lethality of infection dose

Bacterial cultures of *E. coli* ATCC 25922 and *S. aureus* ATCC 43300 were prepared in LB broth and after establishing colony calibration curves (CFU/mL vs. OD600) to ensure accurate dosing (**Figure S2**), 10 µL of various bacterial concentrations at logarithmic growth phase were injected into the last left proleg of the larvae (10 larvae per group) and monitored for 5 days to determine the lethal dose for each bacterium. The lethal doses found to be $5\times10^6$ CFU/mL for *E. coli* (**Figure S3**) and $5\times10^8$ CFU/mL for *S. aureus* (**Figure S4**).

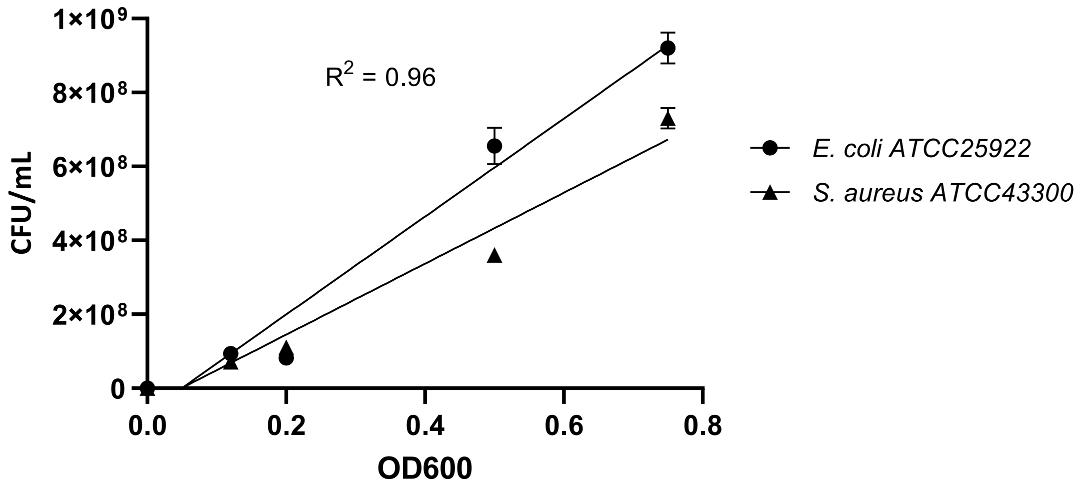


**Figure S2**. Standard curve representing *S. aureus* and *E. coli* (in colony forming units per mL of Lb broth) against absorbance at 600 nanometres. Bacteria were grown in LB broth and plated on LB agar plate according to the drop plate method and incubated at 35±2°C for 24 hours. Experiments were performed in triplicate.

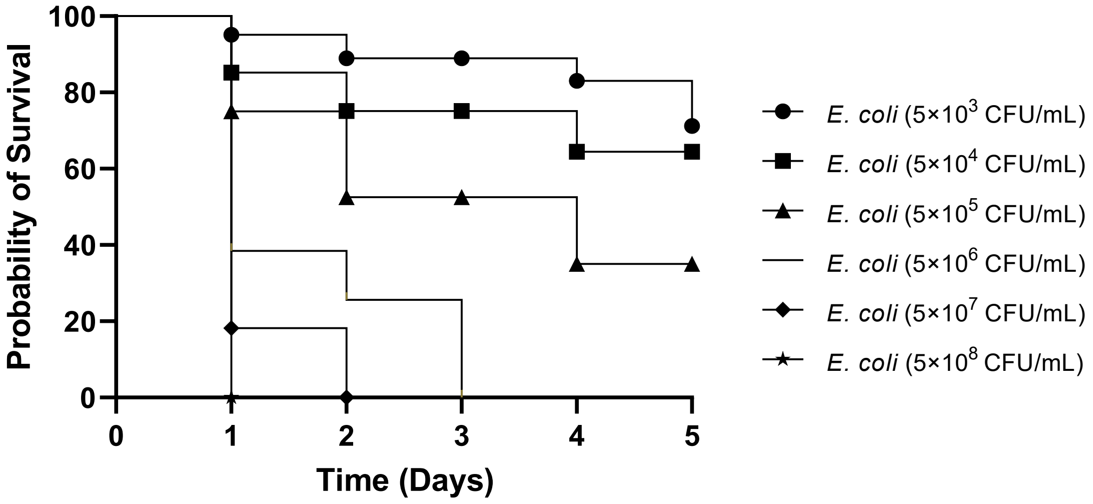


**Figure S3.** Survival Probability of *G. mellonella* larvae after 5 days of infection with *E. coli* ATCC25922 by injecting from $5×10^3$CFU/mL to $5×10^8$ CFU/mL of *the* strain and incubating at 37°C. The number of surviving larvae was recorded every day for a total of 5 days. Statistical analyses were conducted using the log-rank test, with a significance level set at ($p < 0.05$) for multiple comparisons. All experiments were performed in triplicate, with a sample size of ($n = 30$).

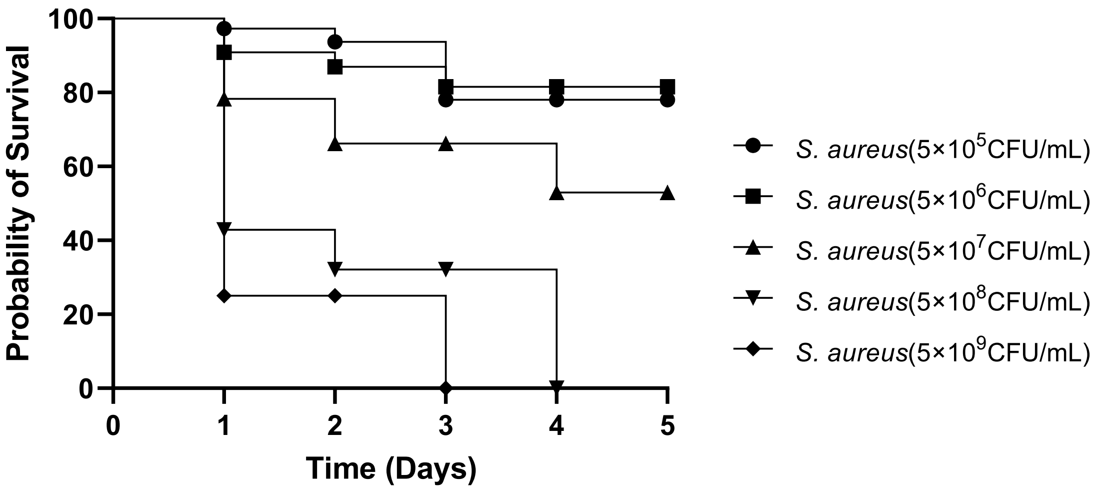


**Figure S4.** Survival Probability of *G. mellonella* larvae after 5 days of infection with *S. aureus* ATCC 43300 by injecting from $5×10^5$CFU/mL to $5×10^9$ CFU/mL of the strain and incubating at 37°C. The number of surviving larvae was recorded every day for a total of 5 days. Statistical analyses were conducted using the log-rank test, with a significance level set at ($p < 0.05$) for multiple comparisons. All experiments were performed in triplicate, with a sample size of ($n = 30$).
